# Transport mechanism of the neuronal excitatory amino acid transporter

**DOI:** 10.1101/2020.06.01.127704

**Authors:** Biao Qiu, Doreen Matthies, Eva Fortea, Zhiheng Yu, Olga Boudker

## Abstract

Human excitatory amino acid transporter 3 (hEAAT3) mediates glutamate uptake in neurons, intestine, and kidney. Here, we report Cryo-EM structures of hEAAT3 in several functional states where the transporter is empty, bound to coupled sodium ions only, or fully loaded with three sodium ions, a proton, and the substrate aspartate. The structures suggest that hEAAT3 operates by an elevator mechanism involving three functionally independent subunits. When the substrate-binding site is near the cytoplasm, it has a remarkably low affinity for the substrate, perhaps facilitating its release and allowing for the rapid transport turnover. The mechanism of the coupled uptake of the sodium ions and the substrate is conserved across evolutionarily distant families and is augmented by coupling to protons in EAATs. The structures further suggest a mechanism by which conserved glutamate mediates proton symport.

## Introduction

Human excitatory amino acid transporters (hEAATs) pump glutamate into cells against steep concentration gradients by utilizing the pre-established transmembrane gradients of sodium and potassium ions and protons as an energy source^1,2^. hEAAT3 is one of five human EAAT subtypes broadly expressed in neurons throughout the brain. It is the major glutamate and aspartate transporter outside of the central nervous system (CNS), including in kidneys and intestines^3–5^. Mutations in hEAAT3 cause dicarboxylic aminoaciduria, a metabolic disorder that leads to the excessive loss of aspartate and glutamate in the urine^6^. Notably, the disorder can be associated with mental retardation and obsessive-compulsive syndrome^7^. The causative role of EAAT3 loss was confirmed using knockout mice, which also developed dicarboxylic aminoaciduria^8^.

In the mammalian brains, EAAT3 is expressed in the somata and dendrites of both excitatory and inhibitory neurons but excluded from presynaptic membranes. EAAT3 total levels in the brain are approximately 100 times lower than EAAT2^9^, which mediates most of the glutamate uptake following synaptic transmission^10^. Nevertheless, altered EAAT3 expression and plasma membrane trafficking are associated with epilepsy, schizophrenia, hypoxia, and decreased activity, which leads to a susceptibility of neurons to oxidative damage, particularly in epilepsy models^11–14^. The relevance of EAAT3 to oxidative stress resistance might be due to its ability, unique among EAATs, to transport cysteine, which is a rate-limiting intermediate in the biosynthesis of the antioxidant glutathione^15–17^. Thus, activators of hEAAT3 might be useful to boost cysteine uptake into neurons under pathologic conditions. Due to the high concentration of glutamate in neurons, hEAAT3 might also run in reverse during ischemia, spilling excitotoxic levels of glutamate into the extracellular space. Therefore, specific inhibitors of hEAAT3 might be clinically helpful in reducing glutamate-mediated damage following transient ischemic conditions and depolarization of membranes.

A detailed understanding of the hEAAT3 structure and mechanism would be required to enable pharmacologic manipulations. Toward this end, we have determined structures of hEAAT3 using Cryo-EM. We show that the transporter is a homotrimer. Each protomer consists of two domains, a central trimerization scaffold domain and a peripheral transport domain containing the substrate-binding site. The individual hEAAT3 protomers undergo elevator-like transitions between the outward- and inward-facing conformations, in which their transport domains localize closer to the extracellular space and the cytoplasm, respectively. They appear to be independent of each other so that we observe trimers with every possible arrangement of the outward- and inward-facing protomers. Remarkably, when we imaged hEAAT3 in the presence of L-aspartate (L-asp) and sodium (Na^+^) ions, we observe that only the outward-facing protomers were bound to the amino acid, while the inward-facing protomers remained substrate-free. We conclude that the substrate-free transporter has a strong preference for the inward-facing conformation while the outward-facing transporter has a significantly higher affinity for the amino acid. By comparing the structures of the transport domain when free of ligands, bound to Na^+^ ions only and bound to Na^+^ ions, protons, and L-asp, we provide mechanistic insights into symport of the amino acid, Na^+^ and protons in hEAAT3.

## Results

### The elevator mechanism of hEAAT3

We heterologously expressed and purified full-length hEAAT3 with two potential glycosylation sites mutated (N178T and N195T), hEAAT3_g_ (**ED Figure 1a - c**). The wild type and mutant showed L-asp uptake in oocytes with a *K_m_* of 39 ± 7 and 63 ± 11 μM (**Figure 1a**), respectively. The purified, liposome-reconstituted hEAAT3_g_ showed electrogenic L-asp and L-glutamate (L-glu) uptake in solid-supported membrane electrophysiology assays (**ED Figure 1d**). The uptake was dependent on a Na^+^ ion gradient and inhibited by (3S)-3-[[3-[[4-(Trifluoromethyl)benzoyl]amino]phenyl]methoxy]-L-aspartic acid (TFB-TBOA) (**ED Figure 1e**). The *K_m_* values for L-asp and L-glu (242 ± 24 and 883 ± 120 μM, respectively) (**Figure 1b**) were higher than in oocytes and mammalian cells^18–20^. The discrepancies might reflect distinct lipid environments or be due to mixed orientations of the hEAAT3_g_ proteins reconstituted into the supported bilayers. Indeed, hEAAT3_g_ showed a significantly higher *K_m_* of reverse transport^21,22^.

**Figure 1.**
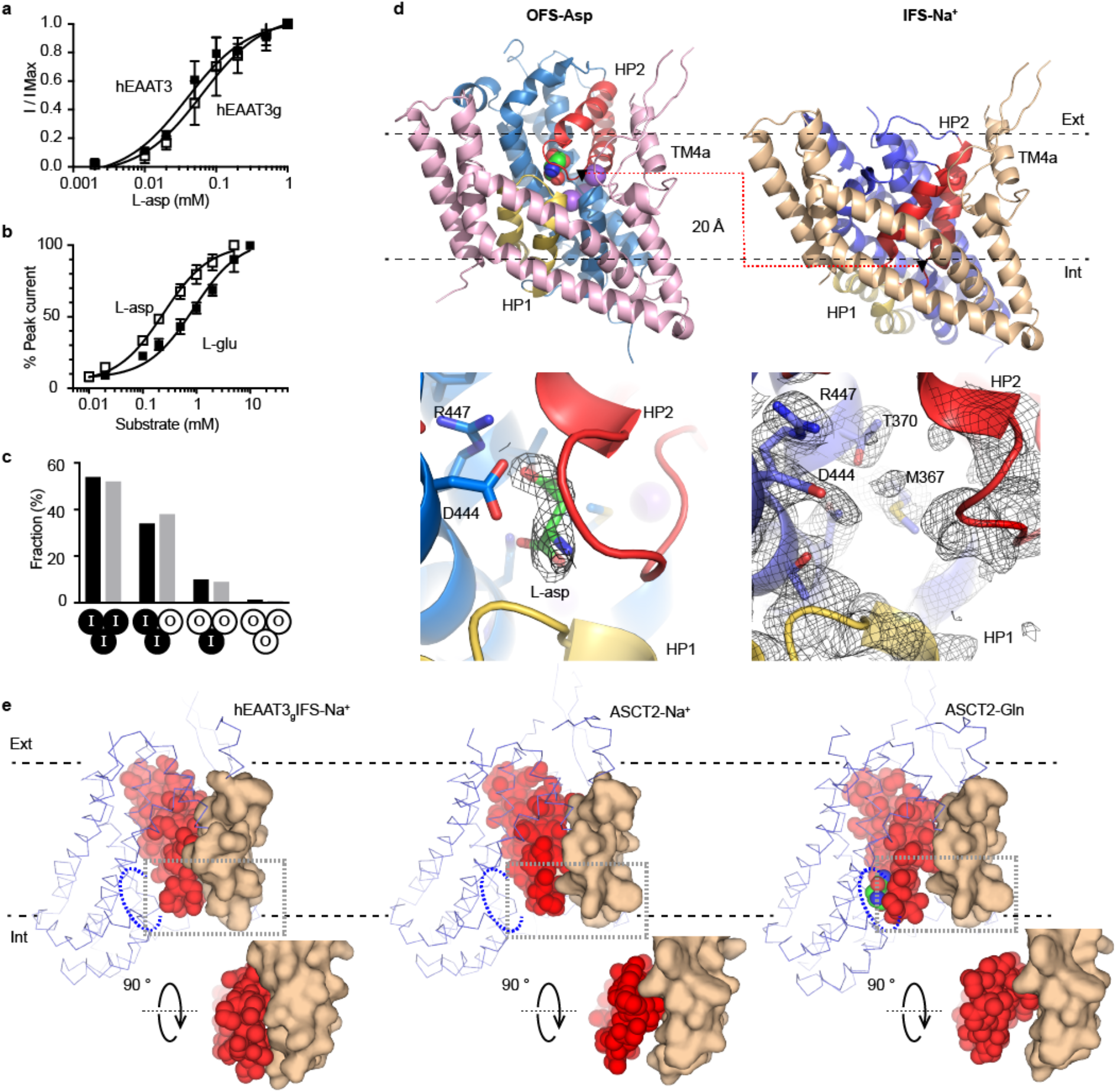
Alternating access mechanism of hEAAT3_g_. **a**, hEAAT3_g_ (open squares) and wild type hEAAT3 (solid squares) show similar L-asp uptake in oocytes. **b**, hEAAT3_g_ reconstituted into solid-supported membranes has a higher apparent affinity for L-asp (open squares) than L-glu (solid squares). **c**, Observed populations of hEAAT3_g_ trimers with different configurations of protomers in the OFS (O) and IFS (I), as indicated below the graph (black columns), are consistent with binomial distributions with OFS probability of 19% (gray columns). The analysis is based on the Cryo-EM imaging data for hEAAT3_g_ in the presence of 200 mM NaCl and 1 mM L-asp. **d**, Structures of single hEAAT3_g_ protomers in the IFS-Na^+^ (left) and OFS-Asp (right) states. The scaffold domains are pink, and beige and the transport domains are in shades of blue. Structurally symmetric HP1 and HP2 are yellow and red, respectively. Substrate and ions are shown as spheres and colored by atom type. Below are the close-up views of the substrate-binding sites. The gray mesh objects are density maps contoured at 5.5 σ for L-asp (left) and 4.5 σ for the protein (right). **e**, Interactions between HP2 (red spheres) and scaffold domain (beige surface) for the inward-facing hEAAT3_g_ bound to Na^+^ ions (left), ASCT2 bound to Na^+^ ions (PDB accession code 6rvx, middle), and ASCT2-Gln bound to glutamine (PDB accession code 6gct, right). Dotted blue ellipses highlight the locations of the substrate-binding site for reference. The rest of the transport domains is shown as blue ribbons. The close-up views are below the panels.

We first acquired Cryo-EM images of hEAAT3_g_ in the presence of 200 mM NaCl and 1 mM L-asp. 2D and 3D classifications were carried out to select a relatively homogenous subset of particles. 3D refinement and reconstruction on the subset with C3 symmetry yielded an electron density map with an overall resolution at 2.85 Å (**Table 1, ED Figure 2a-c and ED Figure 3b**). The trimeric transporter showed an overall architecture (**ED Figure 4a**) similar to those of the homologous archaeal transporters Glt_Ph_ and Glt_Tk_^23–26^, a thermostabilized variant of human EAAT1 (htsEAAT1)^27^, and human neutral amino acid transporter ASCT2^28,29^. The central trimerization scaffold supported three peripheral transport domains in inward-facing states. Consequent symmetry expansion revealed that ~19% of the protomers were in an outward-facing conformation (**ED Figure 2, 3a**). By sorting particle images of 554,920 EAAT3 trimers based on the conformations of their protomers, we estimated the fractions containing three, two, one, and zero inward-facing protomers, which were in good agreement with the binomial distribution (**Figure 1c**). Thus, transport domains sample the outward- and inward-facing orientations independently of each other, consistent with the body of literature^30–32^.

**Table 1:** Cryo-EM data collection, refinement and validation statistics.

**Tabel 1: Cryo-EM data collection, refinement and validation statistics**
|  | IFS-Na<sup>+</sup><br>(EMDB-22011)<br>(PDB-6X2L) | OFS-Asp<br>(EMDB-22014)<br>(PDB-6X2Z) | Asymmetric, 2i1o<br>(EMDB-22020)<br>(PDB-6X3E) | In-Apo<br>(EMDB-22021)<br>(PDB-6X3F) |
| --- | --- | --- | --- | --- |
| **Data collection and processing** |  |  |  |  |
| Magnification | 105,000 | 105,000 | 105,000 | 105,000 |
| Voltage (kV) | 300 | 300 | 300 | 300 |
| Electron exposure (e-/Å<sup>2</sup>) | 60 | 60 | 60 | 60 |
| Defocus range (μm) | -0.5 to -1.5 | -0.5 to -1.5 | -0.5 to -1.5 | -0.5 to -1.5 |
| Pixel size (Å) | 0.832 | 0.832 | 0.832 | 0.832 |
| Symmetry imposed | C3 | C1 | C1 | C3 |
| Initial particle images (no.) | 1,717,667 | 1,664,760(C3 expand) | 190,599 | 1,918,723 |
| Final particle images (no.) | 554,920 | 312,743 protomer | 158,349 | 435,398 |
| Map resolution (Å) | 2.85 | 3.03 | 3.42 | 3.03 |
| FSC threshold | 0.143 | 0.143 | 0.143 | 0.143 |
| Map resolution range (Å) | 23.96-2.66 | 50-2.66 | 34.24-3.15 | 23.96-2.89 |
| **Refinement** |  |  |  |  |
| Initial model used (PDB code) | ab-initio | IFS-Na<sup>+</sup> | IFS-Na<sup>+</sup> and<br>OFS-Asp | IFS-Na<sup>+</sup> |
| Model resolution (Å) | 2.88 | 3.36 | 3.45 | 3.15 |
| FSC threshold | 0.5 | 0.5 | 0.5 | 0.5 |
| Model resolution range (Å) | 29.17-2.88 | 27.67-3.35 | 27.28-3.45 | 25.65-3.15 |
| Map sharpening *B* factor (Å<sup>2</sup>) | -136.18 | -122.27 | -137.77 | -121.29 |
| **Model composition** |  |  |  |  |
| Non-hydrogen atoms | 9,396 | 3,205 | 9,466 | 9,366 |
| Protein residues | 1,233 | 419 | 1,241 | 1,227 |
| Ligands |  | 4 | 4 |  |
| ***B* factors (Å<sup>2</sup>)** |  |  |  |  |
| Protein | 50.73 | 84.23 | 71.47 | 56.22 |
| Ligand |  | 95.04 | 87.02 | 64.47 |
| **R.m.s. deviations** |  |  |  |  |
| Bond lengths (Å) | 0.009 | 0.005 | 0.006 | 0.006 |
| Bond angles (°) | 0.974 | 0.830 | 0.899 | 0.925 |
| **Validation** |  |  |  |  |
| MolProbity score | 1.16 | 1.60 | 1.27 | 1.51 |
| Clashscore | 2.79 | 6.52 | 3.28 | 3.78 |
| Poor rotamers (%) | 0 | 0 | 0 | 0 |
| **Ramachandran plot** |  |  |  |  |
| Favored (%) | 97.53 | 96.37 | 97.14 | 95.04 |
| Allowed (%) | 2.47 | 3.63 | 2.86 | 4.96 |
| Disallowed (%) | 0 | 0 | 0 | 0 |

**Figure 2.**
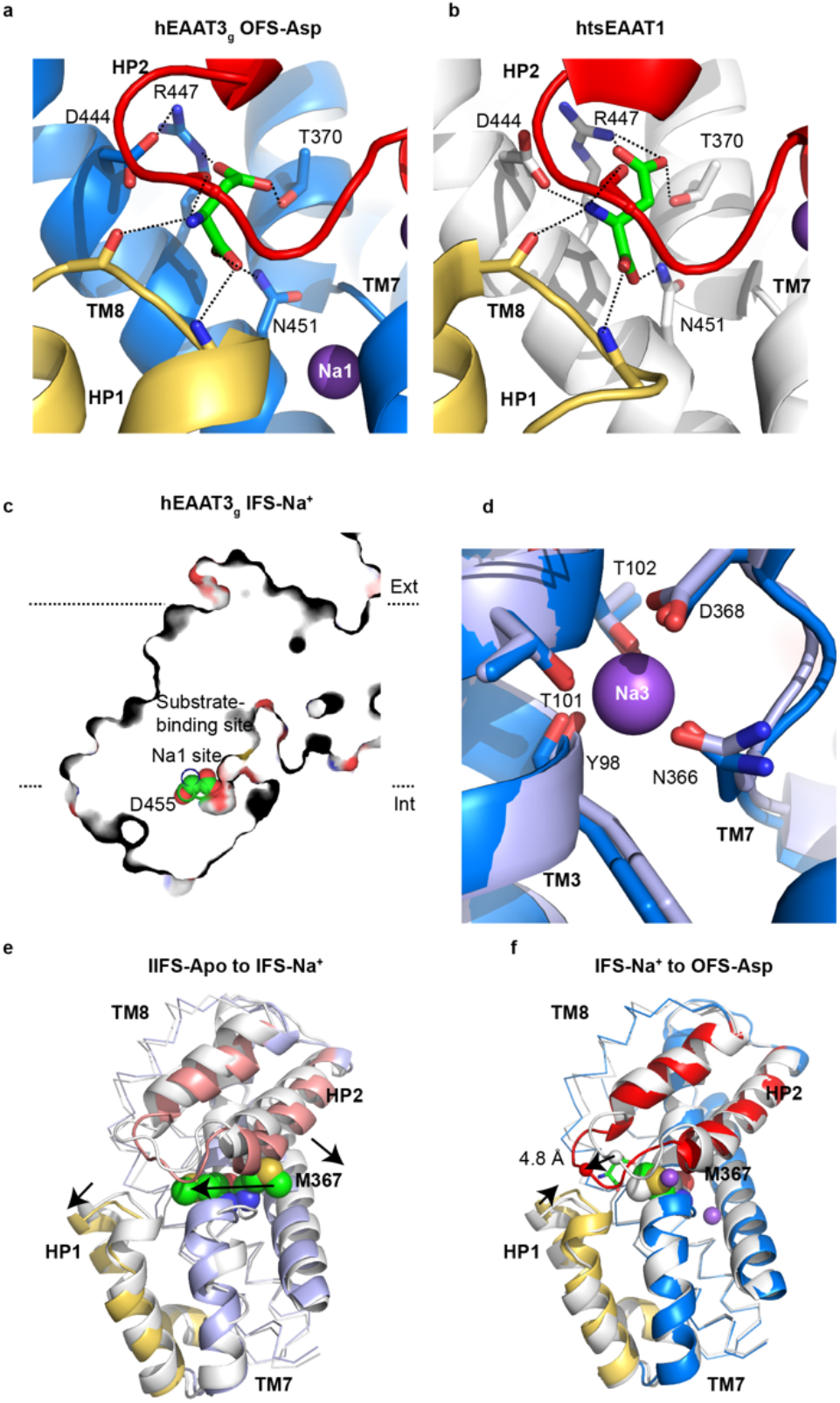
Substrate and sodium coupling in hEAAT3_g_. L-asp binds differently to hEAAT3_g_ (**a**) and thermally stabilized human EAAT1 (htsEAAT1, PDB accession code 5lm4, **b**). Interactions between L-asp and EAATs are shown as dashed lines. HP1 and HP2 are colored yellow and red, respectively, L-asp green, TM 7 and TM 8 blue (hEAAT3_g_) and white (htsEAAT1). **c**, Na1 is solvent-accessible. hEAAT3_g_ is shown as a thin slice taken through the Na1 site. The Na+-coordinating residue D455 is shown as spheres and colored by atom type. **d**, a close-up view of the Na3 site in hEAAT3_g_ OFS-Asp (marine) and IFS-Na^+^ (light blue). **e**, **f**, Conformational changes within the transport domains transitioning from the IFS-Apo to IFS-Na+ state (**e**) and from IFS-Na+ to OFS-Asp state (**f**). The transport domains are white and shades of blue for the starting and ending states, respectively, with HP1 yellow, and HP2 salmon. M367 is shown as spheres and colored by atom type in all states. The arrows indicate conformational changes from IFS-Apo to IFS-Na+ (**e**) and from IFS-Na+ to OFS-Asp (**f**).

**Figure 3.**
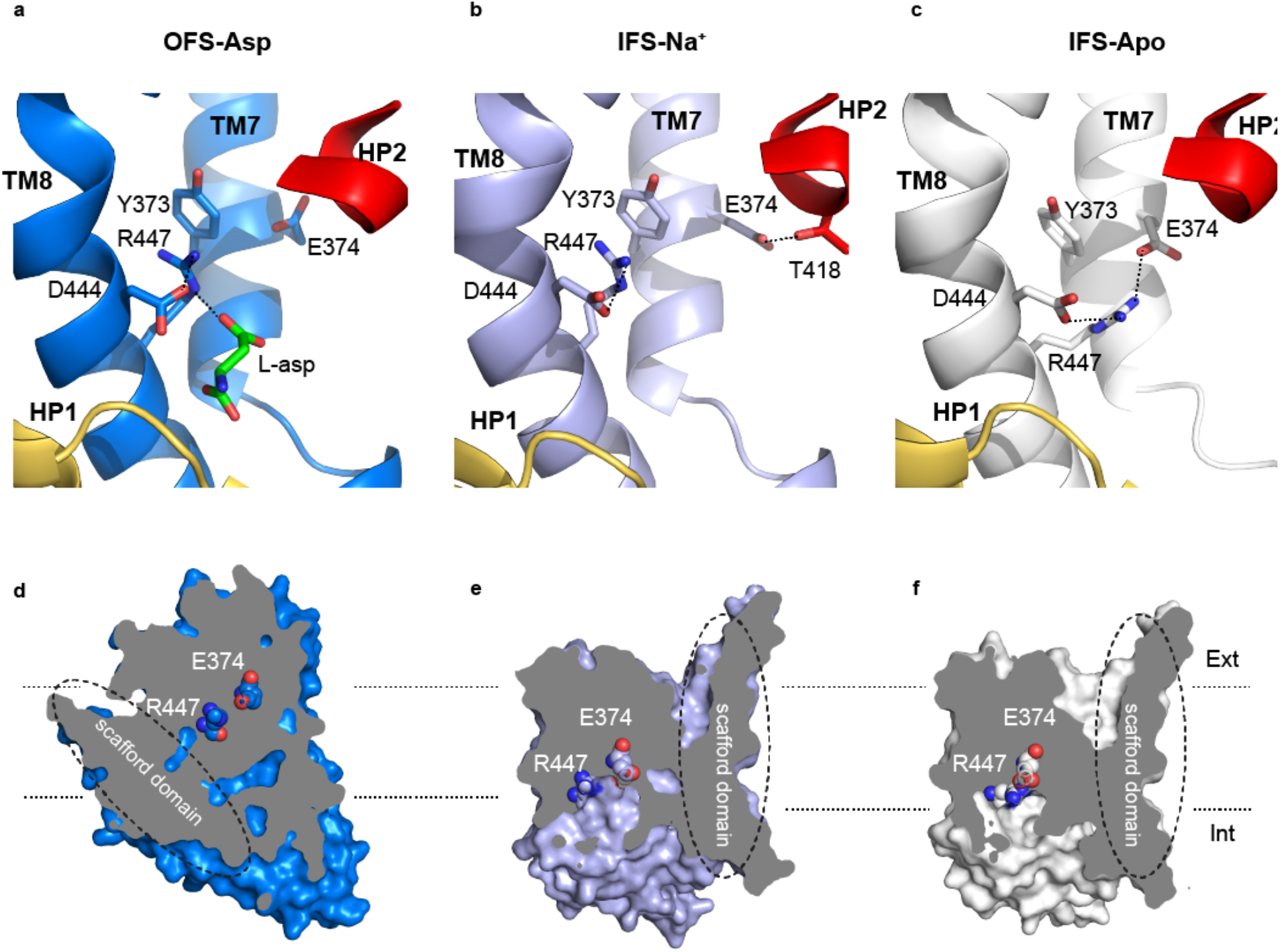
The proton coupling mechanism of hEAAT3_g_. Close-up views of R447 and E374 in the OFS-Asp state (**a**), IFS-Na+ state (**b**), and IFS-Apo state (**c**). Key interactions are shown as dashed lines. HP1 and HP2 are yellow and red, respectively. TMs 7 and 8 are marine, light blue, and white in a, b, and c, respectively. L-asp is green. The position of R447 and E374 relative to the membrane and their solvent accessibility in the OFS-Asp state (**d**), IFS-Na+ (**e**), and IFS-Apo state (**f**). hEAAT3g is shown as a surface sliced approximately through the substrate-binding site and viewed in the membrane plane. Dashed ellipses mark the scaffold domains. R447 and E374 are shown as spheres.

We refined the inward-facing symmetric trimer and the asymmetric trimer with two protomers in the inward-facing state and one protomer in the outward-facing to 3.03 and 3.42 Å resolution, respectively (**Table 1, ED Figure 2d** and **ED Figure 3d**). The map obtained using all particles was virtually indistinguishable from the map based on symmetric inward-facing trimers only. However, it reached a higher resolution, and we used it for further protein model building. We also refined 7,737 symmetric outward-facing trimers and the single outward-facing protomer to 3.69 Å and 3.03 Å resolution, respectively (**Table 1, ED Figures 2e and ED Figure 3a, c**).

In the outward- and inward-facing protomers, the scaffold domain remained mostly unchanged. In contrast, the transport domain moved by ~20 Å from an extracellular to an intracellular position enabled by the inter-domain hinges (**Figure 1d**). Most remarkably, the outward-facing protomers were bound to L-asp (OFS-Asp), while the inward-facing protomers were not (**Figure 1d**). In the inward-facing protomers (IFS-Na^+^), Na^+^ ions are likely bound in at least one of the sites (discussed below), and we observe a characteristic opening of the helical hairpin 2 (HP2) proposed to gate the substrate-binding site^33^. When we imaged hEAAT3_g_ in 20 mM L-asp, the population of the OFS-Asp protomers increased to ~62%, but the inward-facing protomers remained mostly unbound (**ED Figure 5a, ED Figure 7, and Table 2**). Finally, in 10 mM L-glu, we observed only substrate-free IFS-Na^+^ protomers (**ED Figure 5b, ED Figure 7, and Table 2**).

**Table 2:** Cryo-EM data collection and processing.

|  | IFS in 10 mM L-Glu<br>(EMDB-22022) | IFS in 250 mM KCl<br>(EMDB-22023) | Asymmetric 2o1i<br>in 20 mM L-Asp<br>(EMDB-22024) |
| --- | --- | --- | --- |
| **Data collection and processing** |  |  |  |
| Magnification | 105,000 | 105,000 | 105,000 |
| Voltage (kV) | 300 | 300 | 300 |
| Electron exposure (e-/Å<sup>2</sup>) | 60 | 60 | 67.6 |
| Defocus range (µm) | -0.5 to -1.5 | -0.5 to -1.5 | -1 to -2 |
| Pixel size (Å) | 0.832 | 0.832 | 1.096 |
| Symmetry imposed | C3 | C3 | C1 |
| Initial particle images (no.) | 2,884,157 | 1,769,994 | 1,533,624 |
| Final particle images (no.) | 647,481 | 668,093 | 168,777 |
| Map resolution (Å) | 2.96 | 2.85 | 3.71 |
| FSC threshold | 0.143 | 0.143 | 0.143 |
| Map resolution range (Å) | 18.43-2.89 | 18.43-2.69 | 24.11-3.40 |

These results suggest that in the absence of transmembrane electrochemical gradients and substrate, hEAAT3_g_ strongly favors the inward-facing conformation, similar to the archaeal homologue Glt_Tk_^34^. In this state, at least under our imaging conditions, the transporter has a remarkably low substrate affinity, perhaps in tens of mM. Comparing inward-facing hEAAT3_g_ to ASCT2, we observed that in substrate-free proteins, the transport domain leans away from the scaffold, while HP2 remains in contact with the scaffold (**Figure 1e**). Thus, the binding site is open to the cytoplasm. Upon substrate binding to ASCT2, the tip of HP2 closes over the binding pocket. We expect that substrate binding and occlusion in hEAAT3_g_ occur by a similar mechanism. However, the HP2 tip is more extensively engaged with the scaffold in substrate-free hEAAT3_g_ than in ASCT2^28,33^ (**Figure 1e**), perhaps explaining, at least in part, its low affinity for the substrate.

### Distinct mode of substrate-binding to hEAAT3

The substrate-binding site geometry in hEAAT3_g_ OFS-Asp resembles that of the archaeal homologs and htsEAAT1, but L-asp side-chain takes a different rotamer and is coordinated differently in hEAAT3_g_ (**Figure 2a, b, ED Figure 5c**). The main-chain carboxylate of L-asp is coordinated principally by N451_401_ (here and elsewhere, the corresponding residue number in Glt_Ph_ is shown as the subscript for reference) in the transmembrane segment (TM) 8 and S333_278_ in HP1, similar to other homologs^24,27^. In contrast, the side-chain carboxylate and the amino group of L-asp are coordinated differently. Carboxyl oxygens (OD1 and 2) form hydrogen bonds with R447_397_ in TM8, the main-chain amides of HP2 (OD1), and T370_314_ in TM7 (OD2) in the archaeal transporters and htsEAAT1. In hEAAT3_g_, the carboxyl group and R447_397_ face away from each other somewhat. While still hydrogen-bonded by the guanidinium group, the carboxylate appears to be more engaged by T370_314_ (**Figure 2a**). L-asp amino group in other homologs is coordinated primarily by the highly conserved D444_394_ in TM8, but in hEAAT3_g,_ it is coordinated only by the carbonyl oxygens in HP1 and HP2. D444_394_ does not interact with L-asp and, instead, interacts with R447_397_. The structural basis of why the substrate binds differently to hEAAT3_g_ remains unclear because the residues forming the binding site are highly conserved and essential for transport^20,35,36^. Reduced reliance on R447_397_ and coordination of the arginine by D444_394_ might explain why hEAAT3 accepts cysteine as a substrate while the other homologs do not.

### Na^+^ and substrate coupling

We observed density corresponding to Na^+^ ions at all three conserved Na^+^-binding sites Na1, 2, and 3 in the OFS-Asp protomer (**ED Figure 5d**). In the IFS-Na^+^ protomers, L-asp and Na2 sites are distorted and empty because of the open conformation of HP2. The coordinating groups of the Na1 site, located below the substrate-binding site, are in place, but we did not observe any density for the ion. Moreover, it appears that the site, in particular, the critical D455_405_ residue, is solvent-accessible (**Figure 2c**). Although the density for Na^+^ ion in the Na3 site is unresolved, all coordinating residues are in place, and the site is likely occupied (**Figure 2d**). These results suggest that the Na3 site has the highest affinity for the ions, and might be the first to bind, as previously suggested^26^. In contrast, the Na1 site might remain only partially occupied at 200 mM Na^+^ ions used during grid preparation and might be in rapid exchange with the bulk solution.

To explore the structural underpinning of the coupled substrate and Na^+^ ion symport, we imaged hEAAT3_g_ in the absence of Na^+^ ions and amino acids in buffer containing 100 mM choline chloride (apo state). Following data processing, we only observed inward-facing symmetric trimers (IFS-Apo), which we refined to 3.03 Å resolution (**Table 1, ED Figure 3e**). The overall architecture of IFS-Apo is similar to IFS-Na^+^, but the transport domain swings further away from the scaffold and shifts outward slightly. The transport domains superimposed with an overall r.m.s.d. of 1.3 Å, with the most significant conformational changes occurring in TM7 and HP2 **(Figure 2e)**. In further considering the coupling mechanism, we took advantage of the observations that the conformational changes underlying ion and substrate binding are confined to transport domains and similar in the outward- and inward-facing states of glutamate transporters^34,37,38^. We, therefore, compared the transport domains of IFS-Apo, IFS-Na^+^, and OFS-Asp to visualize structural events during binding.

A critical structural change upon binding of Na^+^ ions to the apo protein is the repositioning of M367_311_, part of the highly conserved NMD motif in TM7. When Na^+^ ions bind to Na1 and Na3 sites, coordinated, in part, by N366_310_, M367_311_ sidechain flips from pointing into the lipid bilayer to pointing toward the substrate-binding site, where it also contributes to the Na2 site **(Figure 2e)**^26^. The flipped-out M367_311_ sidechain, as seen in IFS-Apo, would sterically clash with the HP2 conformation observed in the OFS-Asp, perhaps ensuring that L-asp does not bind in the absence of the ions **(Figure 2e, f)**. Once Na^+^ ions bind to Na1 and Na3 sites, M367_311_ and HP2 are repositioned in a manner compatible with substrate binding, which causes only minimal further structural changes, limited to the closure of the HP2 tip and the formation of the Na2 site (**Figure 2f**). Similar movements of M367_311_ contribute to the coupled binding of Na^+^ ions and L-asp in Glt_Ph_^24,37^, suggesting that the mechanism is conserved in glutamate transporters of vastly different evolutionary origins.

Interestingly, in contrast to apo Glt_Ph_ and Glt_Tk_, in which HP2 can close and occlude the substrate-binding site, HP2 remains open in hEAAT3_g_. Closure of HP2 is required for the rapid return of the transport domain into the outward-facing state. In human EAATs, coupled to potassium counter-transport, potassium binding is likely necessary to close HP2. We speculate that the inability of hEAAT3_g_ IFS-Apo to close HP2 is related to the position of the R447_397_ sidechain. In the apo archaeal transporters, R447_397_ moves into the substrate-binding site where it can form direct or through-water hydrogen bonds with the main-chain carbonyl oxygens of the HP2 tip (**ED Figure 6**). In hEAAT3_g_ IFS-Apo, R447_397_ is engaged by a TM7 residue E374_318_ (glutamine in archaeal homologs) and is not available to close HP2. Unfortunately, we were unable to visualize potassium binding to hEAAT3_g_ because the Cryo-EM structure obtained in the presence of 250 mM KCl looked identical to IFS-Apo (not shown).

### Gating coupled to proton binding

The working cycle of EAATs involves the symport of a proton along with the amino acid and Na^+^ ions. E374_318_ in EAAT3 and the equivalent residues in other EAATs might be the proton acceptors, and mutating them to glutamines resulted in substrate translocation only in an exchange mode^39^. Consistently, in archaeal homologs and ASCT2, which are not proton-coupled, E374_318_ is replaced by a glutamine. Furthermore, mutating R447_397_ in EAAT3 to cysteine abolished aspartate and glutamate transport, but allowed cysteine transport in an exchange mode, suggesting that R447 is also involved in proton coupling^35^. In OFS-Asp hEAAT3_g_, E374_318_ is occluded from the solution by HP2 and makes no interactions with other residues (**Figure 3a, d**). In IFS-Na^+^, HP2 is open, and E374_318_ is positioned at the far end of the cavity, where it forms a hydrogen bond with T418_362_. R447_397_ interacts with D444_394_ and, through a cation-Π interaction, Y373_317_, as it does in the OFS-Asp (**Figure 3b, e**). Finally, in IFS-Apo, the sidechains of R447_397_ and E374_318_ flip towards each other to form a salt bridge (**Figure 3c, f**). The interaction between R447_397_ and Y373_317_ breaks and R447_397_ descends into the substrate-binding site. We calculated the pK_a-s_ of E374_318_ in the apo, Na^+^-bound, and L-asp-bound states using PROPKA^40^. The obtained values of 7.7, 7.5, and 8.5, respectively, were similar to the experimentally measured pK_a-s_ for proton binding to EAAT3^21^. Increased pK_a_ values and occlusion from solvent suggest that E374_318_ is more likely to bind protons as first Na^+^ ions and L-asp bind to the transporter. In this manner, binding of protons, Na^+^ ions, and the substrate are coupled. During the return of the apo transport domain into the outward-facing state, the interactions between E374_318_ and R447_397_ would allow the occlusion of the deprotonated residue.

## Discussion

EAAT3 is the fastest EAAT subtype with turnover times of ~10 msec. Like other members of the family, it operates by an elevator mechanism. The movements of the individual domains within the trimer are independent of each other, leading to a stochastic distribution of the outward- and inward-facing protomers. Interestingly, we find that the affinity of the inward-facing state is remarkably low, such that even at 20 mM L-asp, we only observe a very week substrate-like density. In contrast, the substrate affinity of the outward-facing state must be considerably higher, and all outward-facing protomers showed bound L-asp. In that regard, the hEAAT3_g_ differs from the archaeal Glt_Ph_, which showed comparable L-asp affinities in the outward- and inward-facing states^41^. Thus, our data suggest that during the transport cycle of EAAT3, the inward-facing substrate-bound closed state is a high-energy transient state and that the release of the substrate is rapid. We speculate that also potassium-bound inward-facing closed conformation might be a high-energy state, perhaps explaining why we did not observe it in our imaging experiments. This hypothesis is consistent with the reorientation of the potassium-bound transport domain being the rate-limiting step of the transport cycle^42^.

Collectively, our data, together with the published work, suggest that a two-prong structural mechanism of Na^+^ and amino acid symport is conserved in evolution from archaea to humans. First, binding of Na^+^ ions to the Na1 and Na3 sites is a prerequisite for the binding of substrate and Na^+^ in the Na2 site and leads to movements of two critical residues: R447_397_ out of the binding site to make space for the amino acid, and M367_311_ into the binding site, where it coordinates Na^+^ in the Na2 site. Second, uncoupled transport of Na^+^ ions bound to Na1 and Na2 sites is prevented because the closure of the HP2 tip, essential to allow translocation, requires binding of the substrate and Na^+^ in the Na2 site. In EAATs, the coupling to E374_318_ protonation enhances both steps of the mechanism. Breaking of the interaction between E374_318_ and R447_397_, favored by E374_318_ protonation, is necessary to make the arginine available to coordinate the substrate. Furthermore, the protonation of E374_318_ is likely required to allow HP2 closure, which completely occludes the residues in a non-polar protein interior.

## Author contribution and interest conflict

B.Q. and O.B. designed the experiments; B.Q. performed the experiments, processed and analyzed data, and refined the molecular models; D.M. optimized grid preparation and performed preliminary data processing; D.M and B.Q. collected Cryo-EM data with Z.Y. overseeing all aspects of the microscope operation; B.Q. and E.F. performed electrophysiology experiments; B.Q. and O.B. wrote the manuscript with input from all authors.

## Competing interests

The authors declare no competing financial interests.

## Acknowledgments

We thank the Weill Cornell Medicine mass spectrometry facility for verifying the protein identity and the NYSBC Simons EM center for access to the microscopes. We thank Misha Kopylov at NYSBC Simons EM center for assistance with data collection and Dr. Shengliu Wang from MSKCC for the initial scripts for particle sorting. We thank Dr. Xiaoyu Wang and Didar Ciftci for useful discussions.

## Data availability

Atomic coordinates for the Cryo-EM structures have been deposited in the Protein Data Bank under accession codes 6X2L, 6X2Z, 6X3E, and 6X3F. The corresponding Cryo-EM maps have been deposited in the Electron Microscopy Data Bank (EMDB) under accession codes 22011, 22014, 22020, and 22021, respectively. Additional Cryo-EM maps have been deposited into EMDB under accession codes 22022, 22023, and 22024. The other data that support the findings of this study are available from the corresponding authors upon reasonable request.

## Supplementary Methods

### Protein expression and purification

The codon-optimized full-length human EAAT3 was cloned into a modified pCDNA 3.1 (+) plasmid (*Invitrogen*), which has an N-terminal Strep tag-II followed by an enhanced green fluorescent protein (eGFP) and PreScission protease site. Glycosylation sites, N178 and N195, were mutated to threonine by site-directed mutagenesis. HEK293F cells (*Invitrogen*) at a density of 2.5 × 10^6^ cells /ml cultured in the FreeStyle^TM^293 medium (*Gibco*) were transiently transfected using 3 mg of appropriate plasmid using poly-ethylenimine (PEI) (*Polysciences*) as a 1:3 plasmid to PEI weight ratio. The cells were diluted with an equal volume of fresh FreeStyle^TM^293 medium 6 hours following the transfection, and valproic acid (*Sigma*) was added to a final concentration of 2.2 mM to boost protein expression 12 h later. The cells were collected ~48 hours after transfection by centrifugation at 4,000 g for 10 min at 4°C. Cell pellets were re-suspended in buffer containing in mM 50 Tris-Cl pH 8.0, 1 L-asp, 1 EDTA, 1 TCEP, 1 PMSF, and 1:200 dilution of mammalian protease inhibitor cocktail (*Sigma*). The re-suspended cells were disrupted using EmulsiFlex-C3 cell homogenizer (*Avestin*) or flash-frozen by liquid nitrogen and stored at −80°C for further use. Cell debris was removed by centrifugation at 10,000 g for 15 min at 4°C, and membranes were harvested by ultracentrifugation at 186,000 g for 1 h at 4°C. The membrane pellets were re-suspended and homogenized using a Dounce homogenizer in buffer containing in mM 50 Tris-Cl pH 8.0, 200 NaCl, 1 aspartate, EDTA, 1 TCEP, 1 PMSF, 1:200 dilution of mammalian protease inhibitor cocktail, and 10% (v:v) glycerol. The membranes were incubated with 2% dodecyl-β-D-maltopyranoside (DDM, *Anatrace*) and 0.4% cholesteryl hemisuccinate (CHS, *Sigma*) for 1 h at 4°C. After ultracentrifugation at 186,000 g for 1 h at 4°C to remove insoluble material, the supernatant was incubated with Strep-Tactin sepharose resin (*GE Healthcare*) for 1 h at 4°C. For the L-asp containing samples, the resin was washed by 8 column volumes of buffer containing 50 mM Tris-Cl pH 8.0, 200 mM NaCl, 0.06% Digitonin.glyco-diosgenin (GDN, *Anatrace*), 1 mM TCEP, 5% glycerol (wash buffer), and 1 mM L-asp. Protein was eluted with 4 column volumes of the wash buffer supplemented with 2.5 mM D-desthiobiotin (elution buffer). Strep tag-II and eGFP were removed by incubating the protein with homemade PreScission protease at 100:4 protein to protease ratio at 4°C overnight. EAAT3 was further purified by size exclusion chromatography using Superose 6 10/300 column (*GE Healthcare*) pre-equilibrated with 20 mM Tris-Cl pH 8.0, 200 mM NaCl, 1m M TCEP, 0.01% GDN (gel filtration buffer) and either 1 or 20 mM L-asp. To prepare hEAAT3_g_ in the presence of L-glu, L-asp in all buffers was replaced with 10 mM L-glu. The peak fractions were pooled, and the protein was concentrated to 6 mg/ml. To prepare Na^+^/L-asp-free samples, the protein was exchanged during size exclusion chromatography into a buffer containing 20 mM Tris-Cl pH 8.0, 1 mM TCEP, 0.01% GDN, and 250 mM KCl or 100 mM choline chloride.

### EM data acquisition

To prepare grids for Cryo-EM imaging, 3-3.5 μl of EAAT3 at ~6 mg/ml were applied to glow-discharged QF R1.2/1.3 300 mesh gold or 400 mesh copper grids (*Quantifoil*, Großlöbichau, Germany). Grids were either blotted for 3 s at 4°C and 100% set humidity and plunge-frozen into liquid ethane using an FEI Mark IV Vitrobot (*FEI company*, part of *Thermo Fisher Scientific*, Hillsboro, OR) or for 7 s using a Leica EM GP (Leica Microsystems Inc, Buffalo Grove, IL). Grids were screened using an FEI Tecnai F20 TEM operated at 200 kV through a Gatan 626 side-entry cryo holder equipped with a Gatan K2 Summit direct detector (*Gatan, Inc.*, Pleasanton, CA) or an FEI Tecnai Spirit BioTWIN with a TVIPS F416 camera. The four here discussed Cryo-EM datasets were collected at a 300 kV FEI Titan Krios cryo-electron microscope (HHMI Janelia Krios2) using SerialEM^43,44^. The microscope was equipped with a GIF Bioquantum energy filter and a post-GIF K3 camera (*Gatan, Inc.*). A 100 μm C2 aperture, a 100 μm objective aperture, and a 20eV energy slit centered around the zero-loss peak were used during data collection. Dose fractionation (movie) data were collected at a nominal magnification of 105,000x, that is calibrated magnification of 60096x, corresponding to a physical pixel size of 0.832 Å/px (0.416 Å/px in the resultant movie data with a binning of 0.5x selected). With the K3 camera in standard (non-CDS) counted mode, a dose rate of 15 e^−^/px/s was employed, and each movie contains 60 frames with a total accumulated dose of 60 e^−^/Å^2^ on the sample. A nominal defocus range between ~ −0.5 to −1.5 μm was applied. The dataset in the presence of 20 mM L-Asp was collected on a 300 kV FEI Titan Krios (NYSBC Krios2) equipped with a K2 Summit direct electron detector (*Gatan, Inc.*) at a nominal magnification of 105,000x, (calibrated magnification 45,620x) corresponding to a physical pixel size of 1.096 Å/px. Each movie stack containing 50 frames was exposed in counting mode for 10 s (0.2 s per frame) with a total dose of about 67.6 e^−^/Å^2^. A nominal defocus range between −1 to −2 μm was applied during data collection.

### Data processing

For the dataset collected for EAAT3 in the presence of 1 mM L-asp, drift correction was performed using MotionCor2^45^, and the CTF parameters of micrographs were estimated using Gctf^46^. All other steps of image processing were performed in Relion 3.0^47^. 1,717,667 particles were selected from 6,178 micrographs using a box of 288 pixels. The extracted particles were binned by two and subjected to two rounds of 2D classification and one round of 3D classification. The 3D class showing a good secondary structure was selected, and the particles were re-extracted using the original pixel size of 0.832 Å.

After 3D refinement with C3 symmetry and post-processing, the resulting 3D reconstruction from 554,920 particles yielded a map at a 2.85 Å global resolution. The particles after refinement were expanded using C3 symmetry, and an inverted mask generated using the inward-facing protomer was applied for signal subtraction (first signal subtraction). The resulting 1,664,760 protomers were subjected to 3D classification with a mask and without alignment. 325,222 particles in an outward-facing state and 1,339,538 particles in an inward-facing state were selected. The following 3D refinement and post-processing using the outward-facing particles yielded a map at 3.2 Å resolution. An inverted mask generated using the outward-facing protomer was applied to the expanded particles for a second-round signal subtraction. 312,743 particles in an outward-facing conformation were selected after masked 3D classification without alignment. A final map at 3.03 Å resolution was obtained after 3D refinement and postprocessing.

We identified trimers with three outward-facing protomers (3o) and trimers with one outward-facing protomer and two inward-facing protomers (2i1o) by analyzing the coordinates of the protomers generated by the 1st round of signal subtraction and 3D classification using a home-written script. In total, 7,377 and 190,599 trimers were extracted from the raw micrographs, respectively. A final map at 3.69 Å resolution was obtained for the 3o particles after refinement and postprocessing. The first round of refinement of the 2i1o particles yielded a map at 3.99 Å resolution. The refined particles were then subjected to an additional 3D classification. 158,349 particles were selected and further refined, yielding the final 2i1o map at 3.42 Å resolution after postprocessing.

Other data sets were processed in Relion 3.0. Drift correlation was performed using Motioncor2 within Relion 3.0, and CTF parameters of micrographs were estimated using CTFFIND^48^. From the dataset collected for EAAT3 in KCl buffer, 1,769,994 particles were auto-picked and extracted with two-fold binning from 7,036 micrographs. 668,093 particles were selected and re-extracted after two rounds of 2D classification and one round of 3D classification. A final map at 2.85 Å resolution was obtained after 3D refinement, postprocessing, and polishing. For the dataset collected in 10 mM L-glutamate, 2,884,157 particles were auto-picked from 9,780 micrographs, and 647,481 particles were selected and re-extracted as above. After 3D refinement, postprocessing, and polishing, a map at 2.96 Å resolution was obtained. From the dataset collected in choline chloride buffer, 1,918,723 particles were auto-picked from 4,473 micrographs, and 435,398 particles were selected and re-extracted. A map at 3.24 Å resolution was obtained after 3D refinement and post-processing. Further CTF-refinement and post-processing improved the map to 3.03 Å. Symmetry expansion with signal subtraction and 3D classification without alignment was also performed on all datasets as described above. No conformational heterogeneity was found. For the dataset collected in the presence of 20 mM L-Asp, 1,533,624 particles were auto-picked from 3,424 micrographs using a box of 220 pixels. The particles were extracted using the original pixel size of 1.096 Å and subjected to two rounds of 2D classification, and one round of 3D classification. One class showing a good secondary structure (168,777 particles) was selected. Following 3D reconstitution, polishing, and CTF refinement, a map at 3.71 Å resolution was obtained.

### Model building and refinement

Model building was carried out in COOT^49^, and the protein models were refined using Phenix^50^. For cross-validation, the final model was displaced and then refined against the first unfiltered half-map. FSC curves were calculated between the refined model and the first unfiltered half-map, the refined model and the second unfiltered half-map (which was not refined), and the refined model and the summed map. The well overlaid FSC curves of the two unfiltered maps indicated no over-fitting. The structure figures were prepared in Chimera^51^ or Pymol (*DeLano Scientific*).

### Proteoliposome reconstitution

Liposomes were prepared using 5:5:2 (w:w) ratio of 1-palmitoyl-2-oleoyl-sn-glycero-3-phosphocholine (POPC), 1-palmitoyl-2-oleoyl-sn-glycero-3-phosphoethanolamine (POPE, *Avanti Polar Lipids*) and CHS. The lipids in chloroform were dried and rehydrated at 20 mg/ml by 10 freeze-thaw cycles in buffer containing 50 mM HEPES-Tris buffer, pH 7.4. The liposomes were diluted to 4 mg/ml in buffer containing 50 mM HEPES/NaOH, pH 7.4, 200 mM NaCl, 1 mM TCEP and 1mM L-asp and extruded 11 times through 400 nm polycarbonate membranes (*Avanti Polar Lipids*) using a syringe extruder (*Avanti Polar Lipids*) to form unilamellar liposomes. Liposomes were destabilized with DDM at a 1:1.5 (w:w) lipid to the detergent ratio for 15 min at room temperature, and the purified EAAT3 protein was added to the mixture at a 1:10 (w:w) protein to lipid ratio for 30 min at room temperature. The detergent removal was carried out by adding 100 mg/ml Bio-Beads SM-2 (*Bio-Rad*) for 30 min at room temperature followed by 5 additional rounds at 4°C. To exchange the internal buffer, the proteoliposomes were pelleted by ultracentrifugation at 100,000 g for 45 min at 4°C, diluted into the desired buffer, and subjected to two freeze-thaw cycles. Centrifugation and freeze-thaw steps were repeated three times. The proteoliposomes were extruded 11 times through 400 nm polycarbonate membranes for immediate use.

### Solid supported membrane (SSM) assay

The internal proteoliposome resting buffer contained 50 mM HEPES/KOH, pH 7.4, 150 mM KCl, 2 mM MgCl_2_. The sensors for SSM assay were prepared according to the instrument manual, and the transport-coupled current was recorded by a double-solution exchange method^52,53^. Briefly, a non-activating buffer containing 50 mM HEPES/NaOH, pH 7.4, 150 mM NaCl, 2 mM MgCl_2_ was flown through the sensor to build the ionic gradients across the membranes of the proteoliposomes deposited on the sensor. The transport-coupled current was initiated by flowing non-activating buffer supplemented with different concentrations of the substrate (activating buffers). Finally, the sensor was rinsed in a resting buffer to restore the sensor. The L-asp and L-glu affinities hEAAT3 were measured using sensors from three independent proteoliposomes preparations, and the data were fitted by the equation:

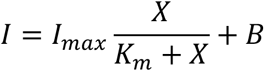

Where *I* is the current measure in the presence of different substrate concentrations, *I_max_* is the fitted maximum current, X is the substrate concentration, and *B* is the baseline offset. TFB-TBOA inhibition was carried out by adding the inhibitor to final concentrations of 10, 3, or 1 μM to non-activating buffer and activating buffer containing 100 μM L-asp. Following recording the transport-activating current, the sensor was restored as described above.

### cRNA preparation, voltage-clamp oocyte recordings, and analysis

Wild type hEAAT3 and hEAAT3_g_ were cloned in a pTLN vector and transcribed *in vitro* using the mMessage mMachine SP6 Kit (*Thermo Fisher Scientific*). *Xenopus laevis* oocytes were purchased from Ecocyte Bio Science (Austin, TX, USA), injected with 25 ng of cRNA, and kept at 18°C in 50% Leibovitz medium, 250 mg/l gentamycin, 1 mM L-glutamine, 10 mM HEPES pH 7.6. Glass microelectrodes were pulled with the resistance of 0.5–3 MΩ and backfilled with 3 M KCl. Currents were recorded 48 h for the wild type and 7 2h for hEAAT3_g_ after injection in ND96 solution (96 mM NaCl, 2 mM KCl, 1.8 mM CaCl_2_, 1 mM MgCl_2_, 5 mM HEPES pH 7.5 using an OC-725C voltage-clamp amplifier (*Warner Instruments*, Hamden, CT). The data were acquired with the Patchmaster (HEKA Elektronik, Lambrecht, Germany) at 5kHz, filtered with Frequency Devices 8-pole Bessel filter at a corner frequency of 2kHz and analyzed using Ana (M. Pusch, Istituto di Biofisica, Genova) and Prism (*GraphPad*). The stimulation protocol for IV curves was as follows: from a holding potential of −30 mV, the voltage was stepped to a variable voltage in 20 mV incremental jumps from −100 mV to +60 mV for 200 ms and stepped back to −30 mV. For calculating *K_m_,* a −100 mV pulse was applied for 200 ms at different ligand concentrations. Each concentration point was proceeded by 0 mM L-asp to control for changes in oocyte leak currents and followed by 1 mM to control for rundown or vice versa. Only oocytes that displayed delta currents bigger than 600 nA in the presence of saturating concentration of ligand were used. Each point in dose-response curves was calculated as follows: (current – leak) / (current at 1 mM L-asp – leak).

### Extended data Figures

**ED Figure 1.**
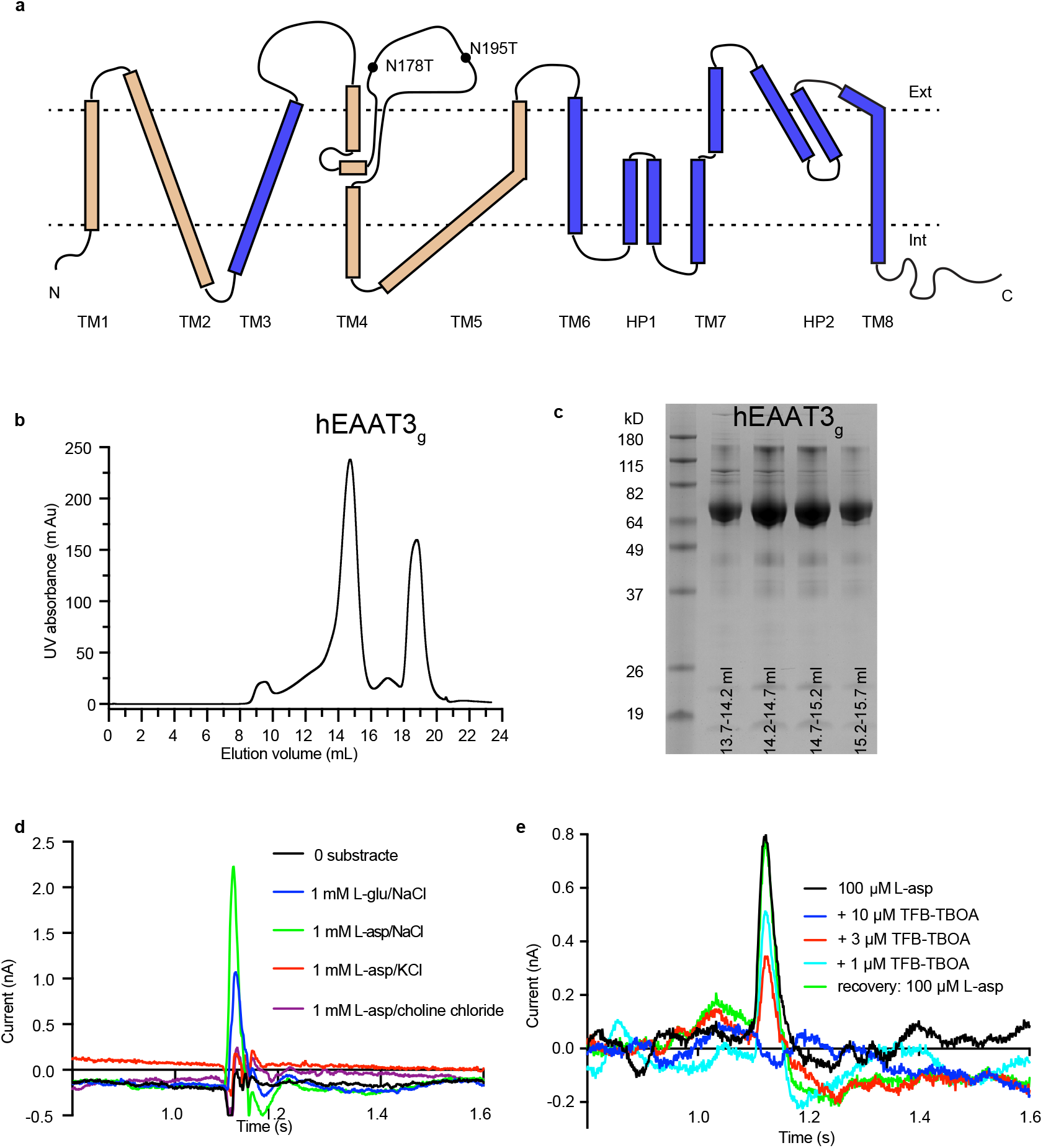
Purification and solid supported membrane assays of EAAT3_g_. **a**, Topology of hEAAT3g with the scaffold domain colored beige, and the transport domain blue. Black circles mark two glycosylation sites, N178 and N195, that were mutated to threonine. **b**, Size exclusion chromatography profile of hEAAT3_g_. **c**, SDS-PAGE analysis of hEAAT3g. **d**, Representative SSM recordings of hEAAT3_g_ in the presence of different ions (green, 1 mM L-asp and 150 mM NaCl; blue, 1 mM glutamate and 150 mM NaCl; red, 1 mM L-asp and 150 mM KCl; purple, 150 mM choline chloride; black, 150 mM NaCl). The substrate transport currents are observed at 1.1 s. Experiments were repeated on two independently prepared batches of proteoliposomes with similar results. **e**, Inhibition by TFB-TBOA of the transport currents induced by 100 μM L-asp. Concentrations of TFB-TBOA were 10 μM (blue), 3 μM (red), and 1 μM (cyan). The uninhibited current was measured before and after each addition of TFB-TBOA to monitor sensor activity. Only the first and last aspartate-induced currents are shown for clarity.

**ED Figure 2.**
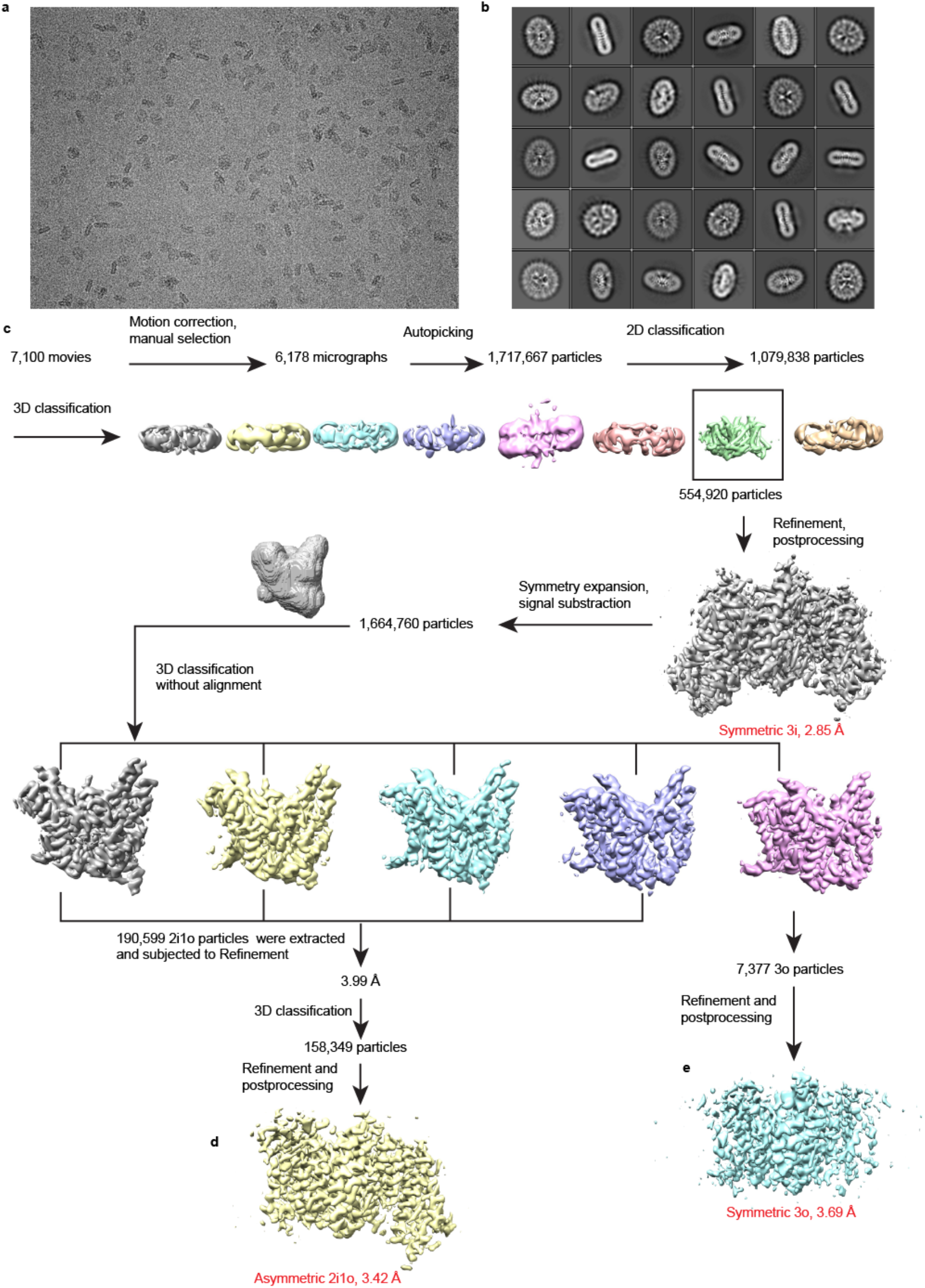
Data processing for hEAAT3_g_ imaged in 200 mM NaCl and 1 mM L-asp. **a,**Example of a micrograph after motion correction. **b**, selected 2D class averages. **c**, Data processing flowchart. Following 3D classification, all particles were refined together using C3 symmetry to yield a map of an inward-facing symmetric trimer, 3i. **d**, Symmetry expansion identified inward- and outward-facing protomers. Trimers containing two inward- and one outward-facing protomer were extracted and refined to yield a map for an asymmetric 2i1o trimer. **e**, Trimers containing all three outward-facing protomers were also extracted and refined using C3 symmetry to yield a symmetric 3o map.

**ED Figure 3.**
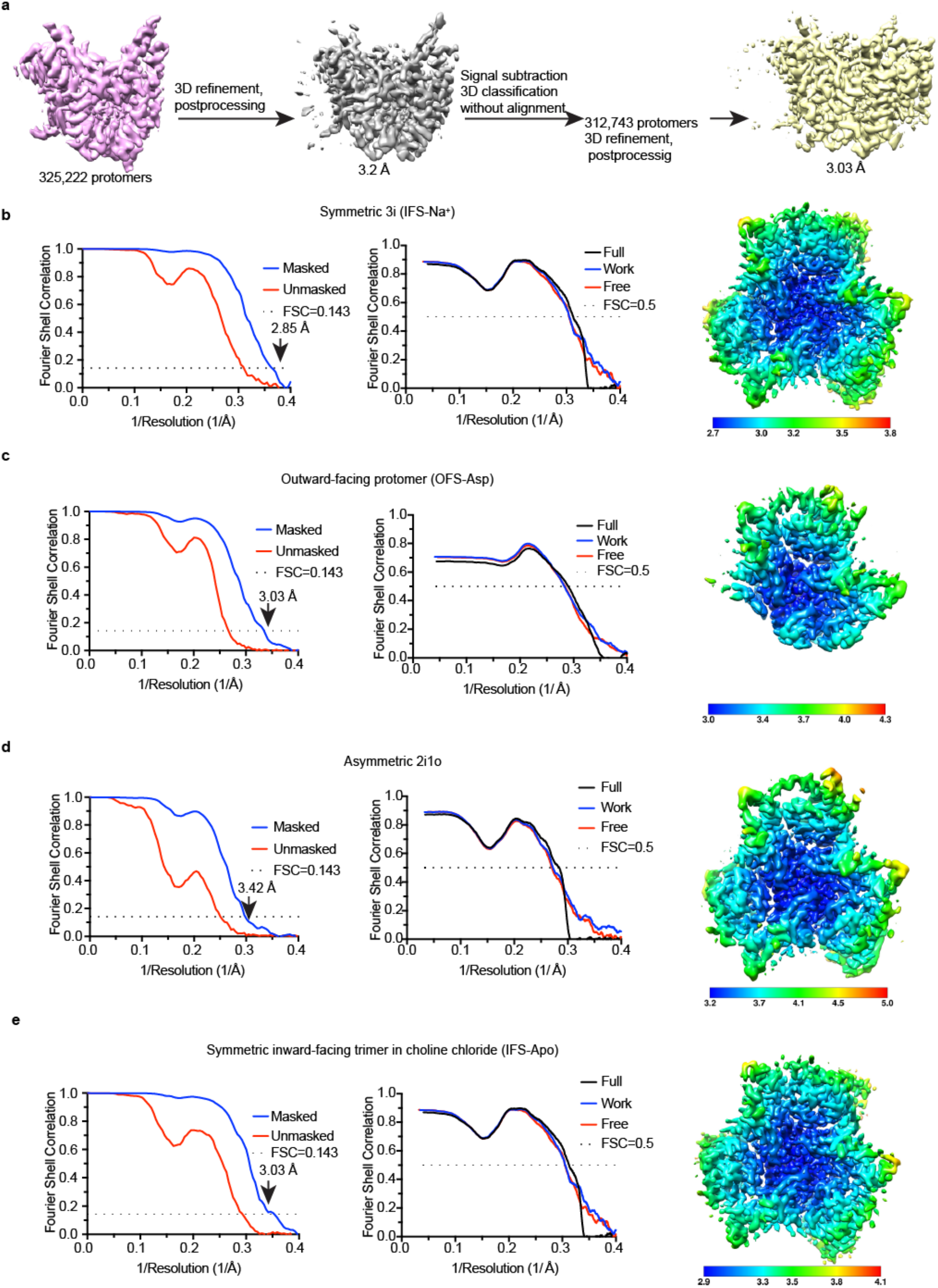
Flowchart of image processing (continued) and validation. **a**, 3D reconstructions of the outward-facing protomer. **b, c, d,** and **e**, Fourier Shell Correlation (FSC) curves for the density maps (left); FSC curves of the refined model versus maps for cross-validation (middle); Density maps colored by the local resolution (right) of symmetric 3i IFS-Na+ (**b**), outward-facing protomer OFS-Asp (**c**), asymmetric 2i1o (**d**), and symmetric inward-facing trimer imaged in choline chloride, IFS-Apo (**e**).

**ED Figure 4.**
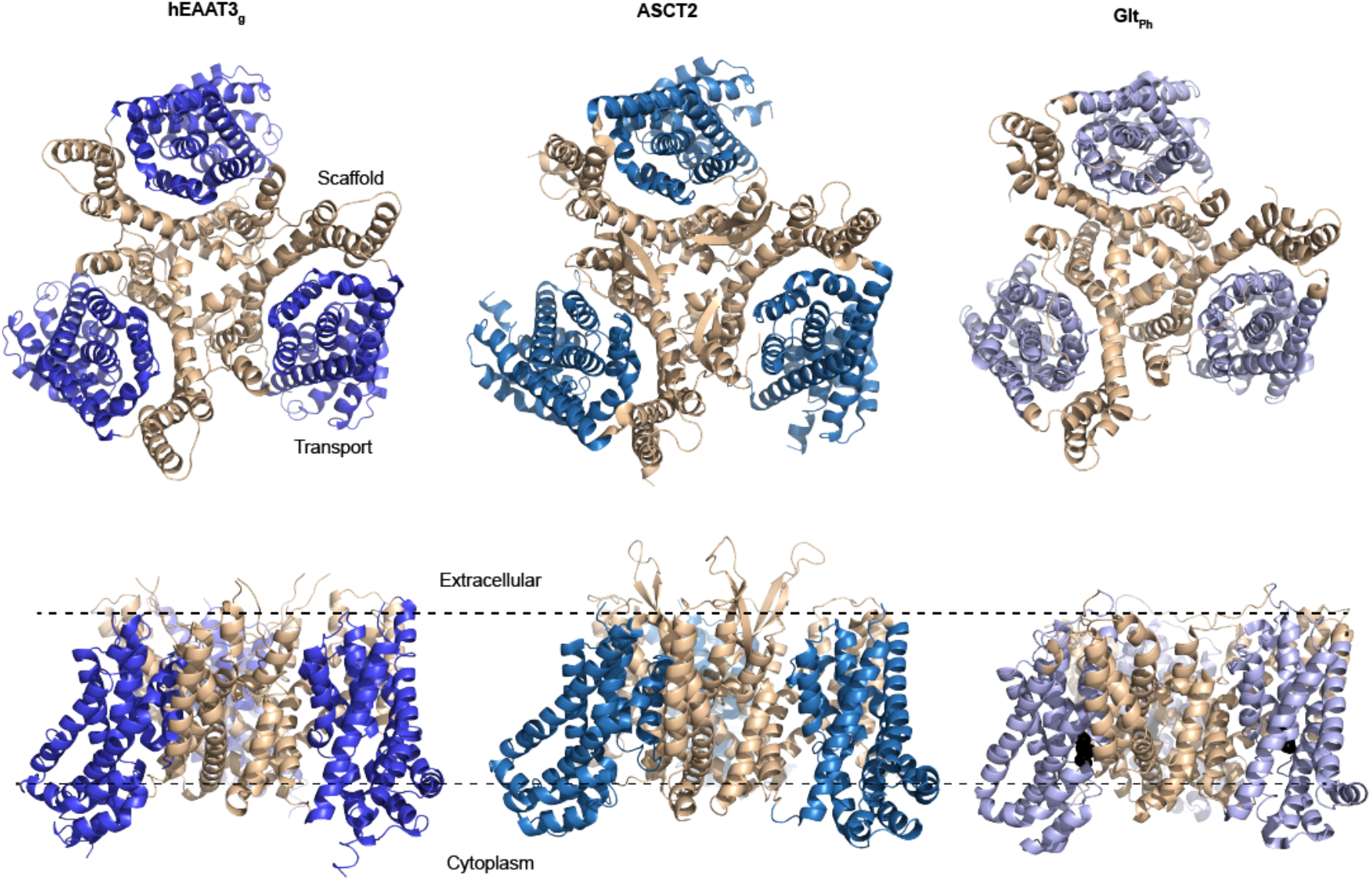
The overall architecture of hEAAT3_g_ resembles those of ASCT2 and Glt_Ph_. Inward-facing transporters are viewed from the extracellular space (top) or in the plane of the membrane (bottom). The transport and scaffold domains are colored beige and shades of blue, respectively. The PDB accession codes are 6rvx for ASCT2 and 3kbc for Glt_Ph_. Black spheres correspond to the bound substrate L-asp in Glt_Ph_.

**ED Figure 5.**
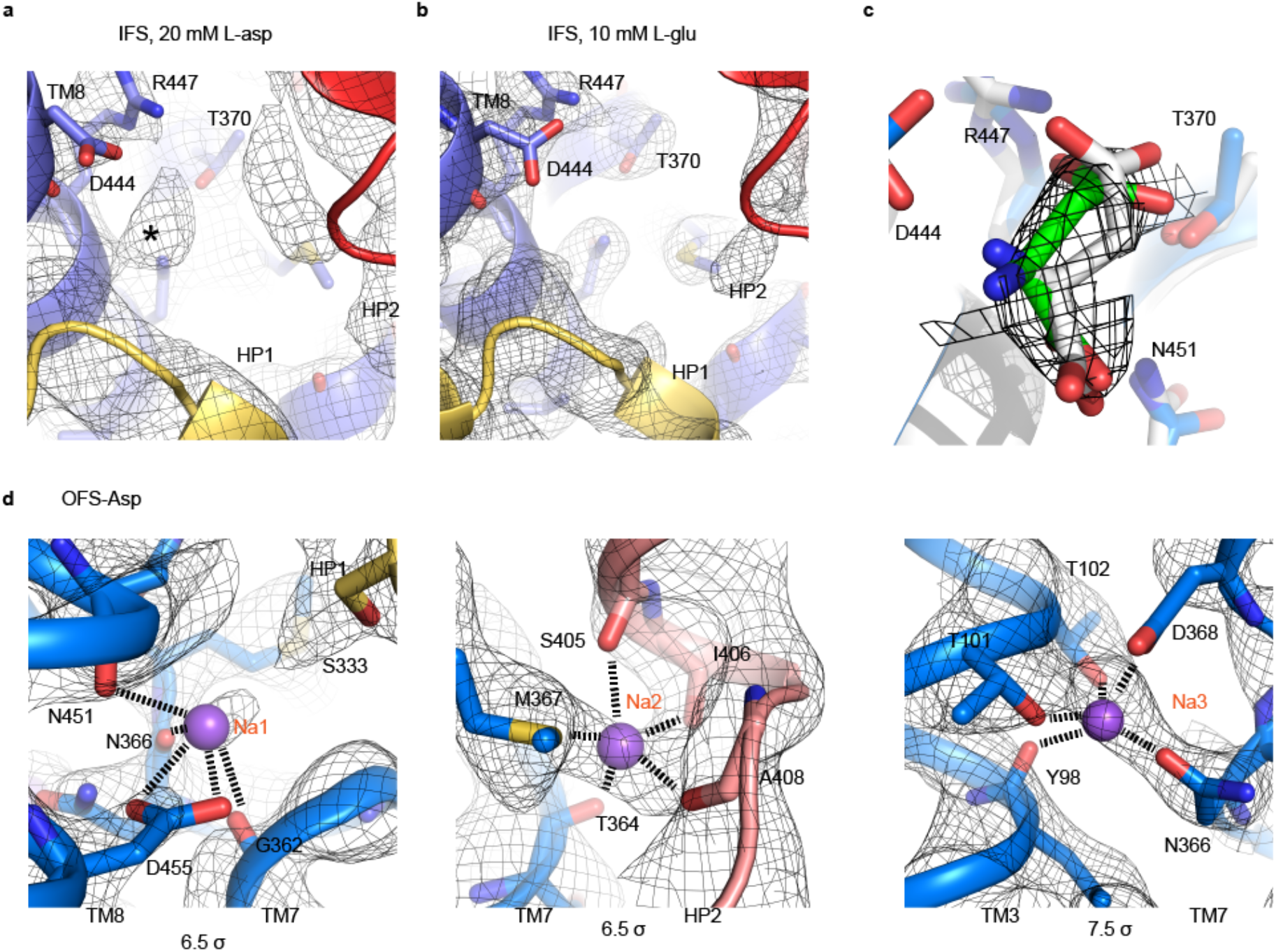
Density maps around the substrate and ion-binding sites. **a** and **b**, Substrate-binding cavity of hEAAT3_g_ in the inward-facing state in the presence of 20 mM L-asp and 10 mM L-glu, respectively. The black mesh shows the density maps. The black star in **a** marks an observed excess density, which might be due to partial occupancy of the substrate. The density around HP2 is fragmented, perhaps reflecting conformational heterogeneity. **c**, Superposition of the modeled L-asp in hEAAT3_g_ (green) and htsEAAT1 (white), showing that the rotamer observed in htsEAAT1 is not compatible with the excess density (black mesh) observed in hEEAT3_g_. **d**, Na^+^-binding sites in the OFS-Asp state of hEAAT3_g_. Na^+^ ions are shown as purple spheres, and their interactions with coordinating atoms are shown as dashed lines. The contour levels of the maps are below the panels.

**ED Figure 6.**
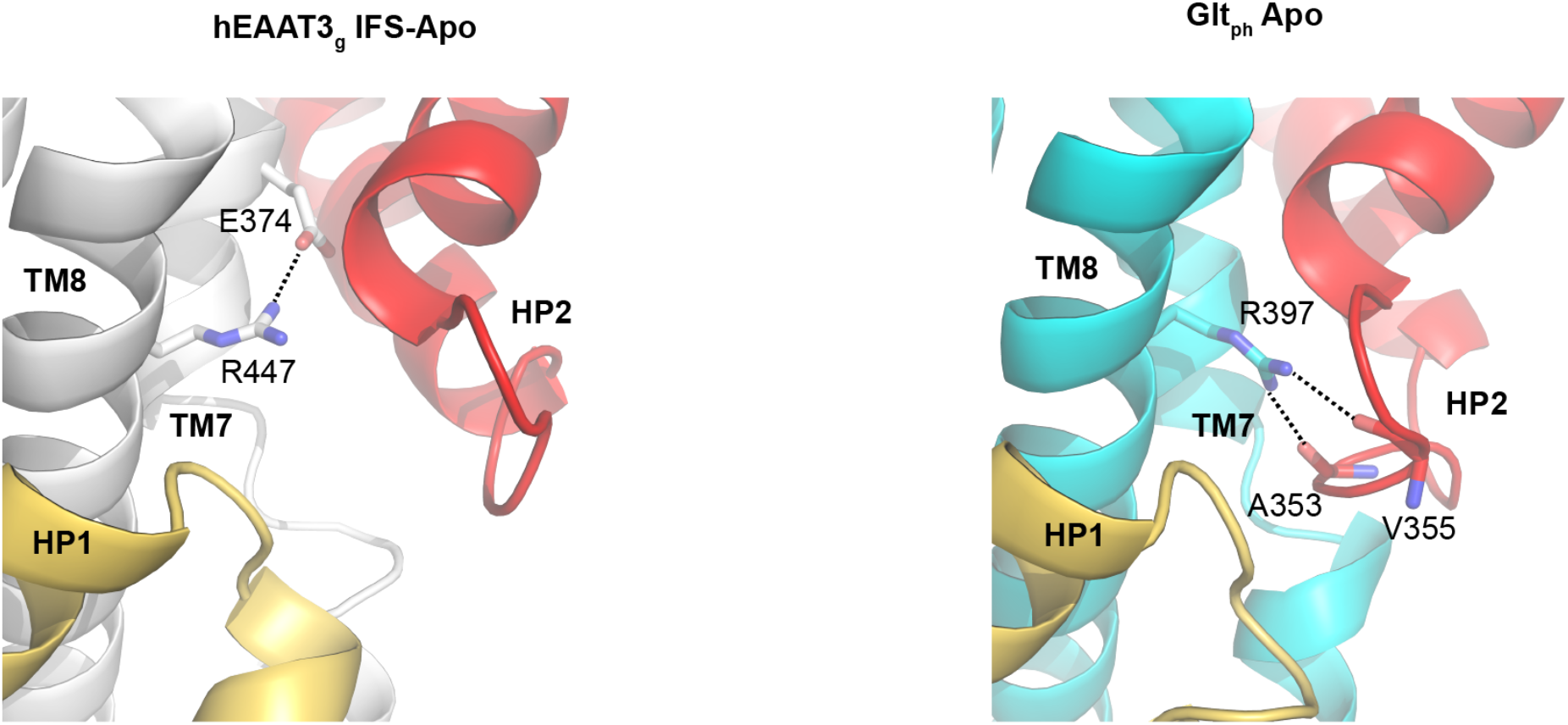
The substrate-binding site of the IFS-Apo hEAAT3_g_ and inward-facing occluded apo Glt_Ph_ (PDB accession code 4p19). Equivalent R447 in hEAAT3_g_ and R397 in Glt_Ph_ are shown as sticks along with their potential interaction partner residues.

**ED Figure 7.**
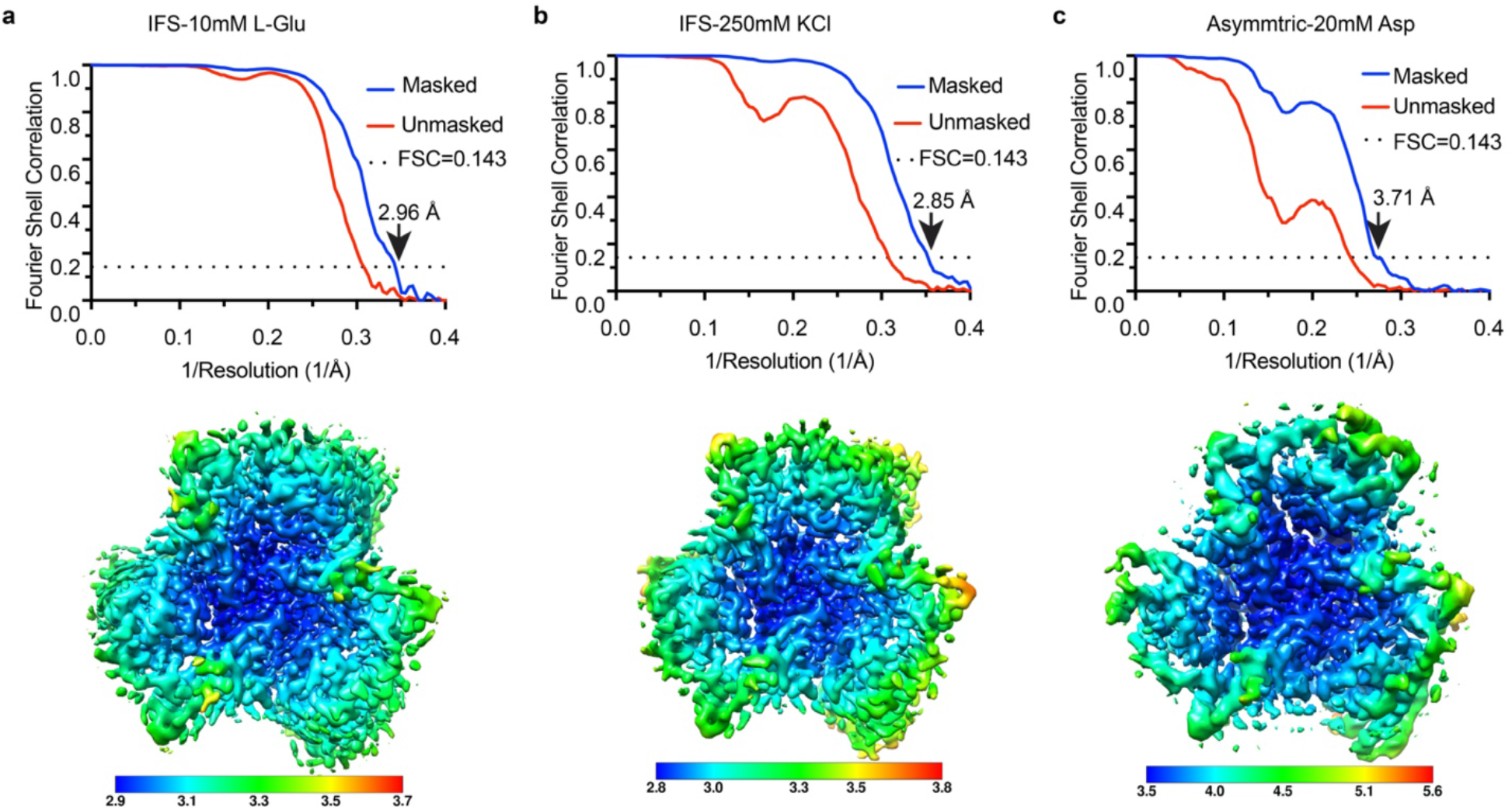
Cryo-EM data processing. Fourier Shell Correlation (FSC) curves for the density maps (top) and density maps colored by the local resolution (bottom) for **a**, IFS in 10mM L-Glu; **b**, IFS in 250mM KCl; **c**, asymmetric trimer, 2o1i, in 2 0mM L-Asp.

## Notes

### Competing Interest Statement

The authors have declared no competing interest.

## References

1 Vandenberg, R. J. & Ryan, R. M. Mechanisms of glutamate transport. Physiological reviews 93, 1621–1657, doi:10.1152/physrev.00007.2013 (2013).

2 Grewer, C., Gameiro, A. & Rauen, T. SLC1 glutamate transporters. Pflugers Archiv : European journal of physiology 466, 3–24, doi:10.1007/s00424-013-1397-7 (2014).

3 Bjorn-Yoshimoto, W. E. & Underhill, S. M. The importance of the excitatory amino acid transporter 3 (EAAT3). Neurochem Int 98, 4–18, doi:10.1016/j.neuint.2016.05.007 (2016).

4 Kanai, Y. & Hediger, M. A. Primary structure and functional characterization of a high-affinity glutamate transporter. Nature 360, 467–471, doi:10.1038/360467a0 (1992).

5 Rothstein, J. D. et al. Localization of neuronal and glial glutamate transporters. Neuron 13, 713–725, doi:10.1016/0896-6273(94)90038-8 (1994).

6 Bailey, C. G. et al. Loss-of-function mutations in the glutamate transporter SLC1A1 cause human dicarboxylic aminoaciduria. J Clin Invest 121, 446–453, doi:10.1172/JCI44474 (2011).

7 Arnold, P. D., Sicard, T., Burroughs, E., Richter, M. A. & Kennedy, J. L. Glutamate transporter gene SLC1A1 associated with obsessive-compulsive disorder. Arch Gen Psychiatry 63, 769–776, doi:10.1001/archpsyc.63.7.769 (2006).

8 Peghini, P., Janzen, J. & Stoffel, W. Glutamate transporter EAAC-1-deficient mice develop dicarboxylic aminoaciduria and behavioral abnormalities but no neurodegeneration. The EMBO journal 16, 3822–3832, doi:10.1093/emboj/16.13.3822 (1997).

9 Holmseth, S. et al. The density of EAAC1 (EAAT3) glutamate transporters expressed by neurons in the mammalian CNS. J Neurosci 32, 6000–6013, doi:10.1523/JNEUROSCI.5347-11.2012 (2012).

10 Rothstein, J. D. et al. Knockout of glutamate transporters reveals a major role for astroglial transport in excitotoxicity and clearance of glutamate. Neuron 16, 675–686, doi:10.1016/s0896-6273(00)80086-0 (1996).

11 Rakhade, S. N. & Loeb, J. A. Focal reduction of neuronal glutamate transporters in human neocortical epilepsy. Epilepsia 49, 226–236, doi:10.1111/j.1528-1167.2007.01310.x (2008).

12 McCullumsmith, R. E. & Meador-Woodruff, J. H. Striatal excitatory amino acid transporter transcript expression in schizophrenia, bipolar disorder, and major depressive disorder. Neuropsychopharmacology 26, 368–375, doi:10.1016/S0893-133X(01)00370-0 (2002).

13 Fujita, H., Sato, K., Wen, T. C., Peng, Y. & Sakanaka, M. Differential expressions of glycine transporter 1 and three glutamate transporter mRNA in the hippocampus of gerbils with transient forebrain ischemia. Journal of cerebral blood flow and metabolism : official journal of the International Society of Cerebral Blood Flow and Metabolism 19, 604–615, doi:10.1097/00004647-199906000-00003 (1999).

14 Bianchi, M. G., Bardelli, D., Chiu, M. & Bussolati, O. Changes in the expression of the glutamate transporter EAAT3/EAAC1 in health and disease. Cell Mol Life Sci 71, 2001–2015, doi:10.1007/s00018-013-1484-0 (2014).

15 Zerangue, N. & Kavanaugh, M. P. Interaction of L-cysteine with a human excitatory amino acid transporter. The Journal of physiology 493 (Pt 2), 419–423, doi:10.1113/jphysiol.1996.sp021393 (1996).

16 Watts, S. D., Torres-Salazar, D., Divito, C. B. & Amara, S. G. Cysteine transport through excitatory amino acid transporter 3 (EAAT3). PloS one 9, e109245, doi:10.1371/journal.pone.0109245 (2014).

17 Aoyama, K. et al. Neuronal glutathione deficiency and age-dependent neurodegeneration in the EAAC1 deficient mouse. Nat Neurosci 9, 119–126, doi:10.1038/nn1609 (2006).

18 Zerangue, N. & Kavanaugh, M. P. Flux coupling in a neuronal glutamate transporter. Nature 383, 634–637, doi:10.1038/383634a0 (1996).

19 Watzke, N., Bamberg, E. & Grewer, C. Early intermediates in the transport cycle of the neuronal excitatory amino acid carrier EAAC1. J Gen Physiol 117, 547–562, doi:10.1085/jgp.117.6.547 (2001).

20 Borre, L. & Kanner, B. I. Coupled, but not uncoupled, fluxes in a neuronal glutamate transporter can be activated by lithium ions. The Journal of biological chemistry 276, 40396–40401, doi:10.1074/jbc.M104926200 (2001).

21 Watzke, N., Rauen, T., Bamberg, E. & Grewer, C. On the mechanism of proton transport by the neuronal excitatory amino acid carrier 1. J Gen Physiol 116, 609–622, doi:10.1085/jgp.116.5.609 (2000).

22 Zhang, Z. et al. Transport direction determines the kinetics of substrate transport by the glutamate transporter EAAC1. Proceedings of the National Academy of Sciences of the United States of America 104, 18025–18030, doi:10.1073/pnas.0704570104 (2007).

23 Yernool, D., Boudker, O., Jin, Y. & Gouaux, E. Structure of a glutamate transporter homologue from Pyrococcus horikoshii. Nature 431, 811–818, doi:10.1038/nature03018 (2004).

24 Boudker, O., Ryan, R. M., Yernool, D., Shimamoto, K. & Gouaux, E. Coupling substrate and ion binding to extracellular gate of a sodium-dependent aspartate transporter. Nature 445, 387–393, doi:10.1038/nature05455 (2007).

25 Reyes, N., Ginter, C. & Boudker, O. Transport mechanism of a bacterial homologue of glutamate transporters. Nature 462, 880–885, doi:10.1038/nature08616 (2009).

26 Guskov, A., Jensen, S., Faustino, I., Marrink, S. J. & Slotboom, D. J. Coupled binding mechanism of three sodium ions and aspartate in the glutamate transporter homologue GltTk. Nat Commun 7, 13420, doi:10.1038/ncomms13420 (2016).

27 Canul-Tec, J. C. et al. Structure and allosteric inhibition of excitatory amino acid transporter 1. Nature 544, 446–451, doi:10.1038/nature22064 (2017).

28 Garaeva, A. A. et al. Cryo-EM structure of the human neutral amino acid transporter ASCT2. Nature structural & molecular biology 25, 515–521, doi:10.1038/s41594-018-0076-y (2018).

29 Yu, X. et al. Cryo-EM structures of the human glutamine transporter SLC1A5 (ASCT2) in the outward-facing conformation. eLife 8, doi:10.7554/eLife.48120 (2019).

30 Grewer, C. et al. Individual subunits of the glutamate transporter EAAC1 homotrimer function independently of each other. Biochemistry 44, 11913–11923, doi:10.1021/bi050987n (2005).

31 Leary, G. P., Stone, E. F., Holley, D. C. & Kavanaugh, M. P. The glutamate and chloride permeation pathways are colocalized in individual neuronal glutamate transporter subunits. J Neurosci 27, 2938–2942, doi:10.1523/JNEUROSCI.4851-06.2007 (2007).

32 Koch, H. P., Brown, R. L. & Larsson, H. P. The glutamate-activated anion conductance in excitatory amino acid transporters is gated independently by the individual subunits. J Neurosci 27, 2943–2947, doi:10.1523/JNEUROSCI.0118-07.2007 (2007).

33 Garaeva, A. A., Guskov, A., Slotboom, D. J. & Paulino, C. A one-gate elevator mechanism for the human neutral amino acid transporter ASCT2. Nat Commun 10, 3427, doi:10.1038/s41467-019-11363-x (2019).

34 Arkhipova, V., Guskov, A. & Slotboom, D. J. Structural ensemble of a glutamate transporter homologue in lipid nanodisc environment. Nat Commun 11, 998, doi:10.1038/s41467-020-14834-8 (2020).

35 Bendahan, A., Armon, A., Madani, N., Kavanaugh, M. P. & Kanner, B. I. Arginine 447 plays a pivotal role in substrate interactions in a neuronal glutamate transporter. The Journal of biological chemistry 275, 37436–37442, doi:10.1074/jbc.M006536200 (2000).

36 Teichman, S., Qu, S. & Kanner, B. I. Conserved asparagine residue located in binding pocket controls cation selectivity and substrate interactions in neuronal glutamate transporter. The Journal of biological chemistry 287, 17198–17205, doi:10.1074/jbc.M112.355040 (2012).

37 Verdon, G., Oh, S., Serio, R. N. & Boudker, O. Coupled ion binding and structural transitions along the transport cycle of glutamate transporters. eLife 3, e02283, doi:10.7554/eLife.02283 (2014).

38 Wang, X. & Boudker, O. Large domain movements through lipid bilayer mediate substrate release and inhibition of glutamate transporters. bioRxiv, 2020.2005.2019.103432, doi:10.1101/2020.05.19.103432 (2020).

39 Grewer, C., Watzke, N., Rauen, T. & Bicho, A. Is the glutamate residue Glu-373 the proton acceptor of the excitatory amino acid carrier 1? The Journal of biological chemistry 278, 2585–2592, doi:10.1074/jbc.M207956200 (2003).

40 Olsson, M. H., Sondergaard, C. R., Rostkowski, M. & Jensen, J. H. PROPKA3: Consistent Treatment of Internal and Surface Residues in Empirical pKa Predictions. J Chem Theory Comput 7, 525–537, doi:10.1021/ct100578z (2011).

41 Akyuz, N., Altman, R. B., Blanchard, S. C. & Boudker, O. Transport dynamics in a glutamate transporter homologue. Nature 502, 114–118, doi:10.1038/nature12265 (2013).

42 Grewer, C., Watzke, N., Wiessner, M. & Rauen, T. Glutamate translocation of the neuronal glutamate transporter EAAC1 occurs within milliseconds. Proceedings of the National Academy of Sciences of the United States of America 97, 9706–9711, doi:10.1073/pnas.160170397 (2000).

43 Mastronarde, D. N. Automated electron microscope tomography using robust prediction of specimen movements. Journal of structural biology 152, 36–51, doi:10.1016/j.jsb.2005.07.007 (2005).

44 Schorb, M., Haberbosch, I., Hagen, W. J. H., Schwab, Y. & Mastronarde, D. N. Software tools for automated transmission electron microscopy. Nature methods 16, 471–477, doi:10.1038/s41592-019-0396-9 (2019).

45 Zheng, S. Q. et al. MotionCor2: anisotropic correction of beam-induced motion for improved cryo-electron microscopy. Nature methods 14, 331–332, doi:10.1038/nmeth.4193 (2017).

46 Zhang, K. Gctf: Real-time CTF determination and correction. Journal of structural biology 193, 1–12, doi:10.1016/j.jsb.2015.11.003 (2016).

47 Scheres, S. H. RELION: implementation of a Bayesian approach to cryo-EM structure determination. Journal of structural biology 180, 519–530, doi:10.1016/j.jsb.2012.09.006 (2012).

48 Rohou, A. & Grigorieff, N. CTFFIND4: Fast and accurate defocus estimation from electron micrographs. Journal of structural biology 192, 216–221, doi:10.1016/j.jsb.2015.08.008 (2015).

49 Emsley, P., Lohkamp, B., Scott, W. G. & Cowtan, K. Features and development of Coot. Acta crystallographica. Section D, Biological crystallography 66, 486–501, doi:10.1107/S0907444910007493 (2010).

50 Adams, P. D. et al. PHENIX: a comprehensive Python-based system for macromolecular structure solution. Acta crystallographica. Section D, Biological crystallography 66, 213–221, doi:10.1107/S0907444909052925 (2010).

51 Pettersen, E. F. et al. UCSF Chimera--a visualization system for exploratory research and analysis. J Comput Chem 25, 1605–1612, doi:10.1002/jcc.20084 (2004).

52 Bazzone, A., Barthmes, M. & Fendler, K. SSM-Based Electrophysiology for Transporter Research. Methods Enzymol 594, 31–83, doi:10.1016/bs.mie.2017.05.008 (2017).

53 Krause, R., Watzke, N., Kelety, B., Dorner, W. & Fendler, K. An automatic electrophysiological assay for the neuronal glutamate transporter mEAAC1. J Neurosci Methods 177, 131–141, doi:10.1016/j.jneumeth.2008.10.005 (2009).

